# Pathway impact analysis (PIS) for robust and comprehensive interpretation of differentially expressed genes (DEGs)

**DOI:** 10.1101/2025.09.08.674792

**Authors:** Yeogha Yoon, Minji An, Hanbi Lee, Wankyu Kim

**Author notes:** To whom correspondence should be addressed. E. mail.

## Abstract

In transcriptome analysis, identifying differentially expressed genes (DEGs) is fundamental for understanding cellular responses and elucidating disease mechanisms. However, conventional DEG selection often relies on arbitrary thresholds for fold change and statistical significance, which are sensitive to experimental noise and sample size, thereby affecting reproducibility and biological interpretability.

We present the Pathway Impact Score (PIS), a data-adaptive method that optimizes DEG thresholds by maximizing cumulative pathway enrichment. Unlike fixed cut-offs, PIS systematically determines dataset-specific thresholds that yield the most coherent and enriched pathways. Validation using the MAQC2 benchmark and 16 dataset of pulmonary fibrosis model suggested that PIS can improve the recovery of true DEGs, while maintaining stability under varying noise levels and replicate counts, and increasing pathway enrichment strength.

PIS also supported high-resolution analysis of dose-dependent transcriptomic responses to phosphodiesterase (PDE) inhibitors, capturing pathway-level perturbations that appeared to align with each compound’s potency. Such analyses may facilitate more precise characterization of drug-induced transcriptional changes, highlighting the potential of PIS for integrative transcriptomic studies.

Overall, PIS provides a practical, biologically informed framework for adaptive DEG selection, particularly suited for meta-analyses and integrative studies where dataset-specific variability needs careful consideration. It is designed for seamless integration into existing transcriptomic analysis pipelines or as a complementary module, requiring minimal additional parameter tuning beyond standard differential expression workflows.

## Introduction

Transcriptome profiling provides a comprehensive view of cellular states and is foundational for downstream analyses such as biomarker discovery and pathway interpretation **[1]**. A critical yet persistent challenge in this field is the reliable identification of differentially expressed genes (DEGs), which traditionally relies on statistical thresholds such as fold change, p-values and false discovery rates (FDR). However, recent critiques—notably by Amrhein *et al*. in *Nature* **[2]** and Benjamin *et al* in *Nature Human Behavior* **[3]**—have challenged the universal acceptance of p < 0.05 as a threshold for ‘statistical significance,’ citing its susceptibility to sample size, data quality, and reproducibility issues. Wasserstein & Lazar (2019) argue for moving ‘beyond p < 0.05’ by adopting more context-aware statistical interpretations **[4]**.

Widely used methods, including DESeq2 **[5]**, edgeR **[6]**, limma **[7]** and others **[8]**, employ these statistical measures to control for variability and multiple testing. These methods typically model count data distributions (e.g., negative binomial in DESeq2 and edgeR) or apply variance modeling (e.g., limma for microarray and RNA-seq with voom transformation). However, threshold selection often depends on arbitrary, user-defined cut-offs, such as fold change > 2 and FDR < 0.05. These criteria, while conventional, may fail to reflect biological context and are highly sensitive to sample size, replicate number, and technical noise **[9–11]**. This lack of standardization not only undermines the reproducibility of DEG-based findings but also limits the biological interpretability of downstream analyses such as pathway enrichment or biomarker discovery **[12]**.

Alternative approaches, including threshold-free methods such as Gene Set Enrichment Analysis (GSEA) **[13]**, Parametric Analysis of Gene Set Enrichment (PAGE) **[14]**, and Generally Applicable Gene-set Enrichment (GAGE) **[15]**, have attempted to circumvent the problem of arbitrary cut-offs by evaluating gene set-level changes across ranked gene lists. While valuable for capturing global pathway perturbations, these methods do not provide a concrete set of DEGs for downstream applications requiring specific gene lists, such as diagnostic panel development or functional validation. Moreover, network-based DEG refinement methods such as MLDEG **[16]**, DEGraph **[17]**, pathDESeq **[18]**, and network propagation algorithms incorporate topological features or machine learning models to improve sensitivity but often introduce additional layers of computational complexity and dependency on network quality, limiting their accessibility for general transcriptome studies.

Here, we present the Pathway Impact Score (PIS), a biologically-informed metric designed to optimize DEG selection by maximizing the total pathway-level perturbation signal. Rather than relying on fixed statistical thresholds, PIS systematically searches for the DEG set that yields the strongest cumulative pathway enrichment, providing a data-adaptive solution that prioritizes biological coherence over arbitrary significance criteria. It relies on the principle that true biological responses often manifest as coordinated changes of functionally related gene sets or pathways **[19**,**20]**, reflecting the evolutionary organization of gene regulatory networks to ensure efficient and robust cellular adaptation across diverse environmental and physiological conditions. By leveraging this property, PIS enables robust and reproducible DEG selection that remains stable across datasets with varying noise levels, replicate numbers, and experimental conditions. Notably, the PIS framework is designed for seamless integration into, or as a complement to, existing transcriptomic analysis pipelines, requiring minimal additional parameter tuning beyond standard differential expression outputs. While demonstrated here using bulk transcriptome datasets, PIS also holds potential for broader applicability to emerging high-resolution technologies such as single-cell and spatial transcriptomics.

## Methods

### Microarray Quality Control 2 (MAQC2) dataset

The benchmark dataset utilized in this study was obtained from the Microarray Quality Control 2 (MAQC2) project ***[21]***, comprising Ambion’s human brain and Stratagene’s human universal reference RNA samples. For analysis, we selected samples with PhiX Control (n = 7; accession SRA010153, NCBI Short-Read Archive). Sequencing reads were aligned to the GRCh38 (v34) reference genome using STAR v2.7.1a ***[22]*** and quantified using featureCounts ***[23]***. DEG analysis was performed using DESeq2 and edgeR with p-values adjusted for multiple testing by the Benjamini-Hochberg method ***[24]***. The MAQC2 project also provides TaqMan qRT-PCR data for 1,044 genes from the same reference samples (GEO accession GSE5350). As a gold standard benchmark, we generated a list of DEGs based on the criteria by Costa-Silva et al. (log2FC > 2, FDR < 0.05) ***[8]***.

### Transcriptome dataset for Mouse pulmonary fibrosis models

Expression datasets from mouse pulmonary fibrosis models were retrieved from the GEO database (Supplementary Table 1). Microarray datasets were pre-processed according to the respective chip manufacturer’s protocols, followed by DEG analysis using limma. For RNA-seq datasets, reads were aligned to the mouse reference genome (mm10) using STAR v2.7.1, quantified with RSEM [25], and analyzed for differential expression using by DESeq2 and edgeR.

### Preparation of non-redundant, standardized pathway dataset

Gene sets from Reactome [26] and Biological Process terms from Gene Ontology [27,28] were compiled to create a standardized reference pathway dataset. Gene set pairs with 90% overlapping genes were considered redundant and merged. Additional redundancies were addressed through manual curation. The curated pathway dataset, comprising 1,071 pathways and covers 15,293 genes, is available at https://github.com/littleheroncodes/PIS.

### Pathway Impact Score (PIS) Analysis

The Pathway Impact Score (PIS) is defined as the total sum of pathway perturbations, calculated by summing the enrichment factors (EFs) across all pathways for a given DEG set. Optimal thresholds and corresponding DEGs are selected by maximizing PIS, evaluated over a series of fold change (FC) and p-value/FDR combinations, or alternatively, across a ranked list of genes.

For a given DEG set A and a pathway gene set B, the enrichment factor (EF) is computed as the ratio of observed to expected overlapping genes, with a pseudocount ps=1 applied for Laplacian smoothing:

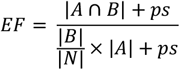

where |N| is the total number of genes.

The *PIS* for the DEG list (A) is calculated as the sum of log-transformed enrichment factors (EFs) for all the pathways with EF ≥ 2:

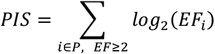

To determine the optimal threshold, a series of DEG sets were generated and PIS was calculated using varying combinations of FC and FDR cut-offs, or by incrementally adding genes from a ranked gene list. The DEG set yielding the highest PIS is selected as optimal.

### Introduction of noises to RNA-seq data

For each vector of read counts (r) corresponding to a gene across replicates, the geometric mean E(r) was calculated. To simulate noise level of k, noise values, n were randomly sampled from the interval:

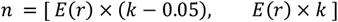

Each sampled noise n was then randomly added to or subtracted from the original read counts to introduce noise. This process was repeated 100 times for each 0.05 increment in the noise level.

### Analysis of drug-perturbed transcriptome on multiple doses

Human Primary Dermal Fibroblast, Neonatal (HDFn) cells (ATCC) were cultured in Fibroblast Basal Medium supplemented with the Fibroblast Growth Kit–Low Serum (ATCC). Cells were seeded onto 96-well plates at densities of 25,000 or 50,000 cells per well and incubated for 24 hours at 37°C in a 5% CO_2_ atmosphere.

Four phosphodiesterase (PDE) inhibitors—roflumilast, rolipram, cilostazol, and milrinone—were administered at 11 concentrations ranging from 7.8 nM to 8,000 nM using a 2-fold serial dilution over 24 hours. Following treatment, cells were lysed with 100 μL of lysis buffer, and total RNA was extracted using the SV 96 Total RNA Isolation System (Promega) according to the manufacturer’s protocol. RNA concentration was quantified using the Quant-iT™ RiboGreen™ RNA Assay Kit (Invitrogen) and stored at −20°C until further use.

RNA sequencing was performed using the DNBSEQ-G400 platform (BGI protocol), and data processing— including read alignment and quantification—followed the same pipeline and reference genome as described for the MAQC2 dataset. Differential gene expression was analyzed, and the results were subjected to PIS analysis, with DEG thresholds optimized within a fold change range of 1.2–2.5 and FDR range of 0.2–0.01.

IC_50_ values for each PDE inhibitor and their respective targets were obtained from the literature and the PubChem bioassay database (Supplementary Table 2) [29]. When multiple IC_50_ values were reported, their mean was used.

## Results

We applied PIS to differential expression results generated by widely used tools—DESeq2 and edgeR—and evaluated PIS against common thresholds (*e*.*g*., fold change and p-value) applied to the same outputs. In this work, threshold-free enrichment methods such as GSEA, PAGE, and GAGE were not employed, as they operate on ranked lists without producing discrete DEGs, making them conceptually distinct from PIS’s objective. Across benchmark and biological datasets, we assessed PIS for its ability to (i) identify dataset-specific thresholds yielding maximal functional enrichment, (ii) remain robust under varying replicate numbers and simulated noise, and (iii) resolve dose-dependent drug perturbation signatures with high biological relevance.

### PIS analysis using MAQC2 dataset

To benchmark the performance of PIS-based DEG selection, we applied the method to the Microarray Quality Control 2 (MAQC2) dataset, using qRT-PCR validation data as a gold standard (Fig. 1A). We hypothesized that the overall biological impact between experimental groups could be captured by the cumulative enrichment of perturbed pathways. Given that genes involved in the same functional processes are frequently co-regulated, we expected biologically meaningful changes to be reflected in coordinated pathway enrichment. This pattern manifests as a peak in the PIS profile, where the number of selected DEGs yields maximal pathway-level enrichment (Fig. 1B). When the DEG set is too small, the signal may be insufficient to support downstream analyses, whereas overly relaxed thresholds may introduce significant noise, obscuring true biological signals. Thus, the PIS peak reflects an optimal balance between signal sensitivity and false discovery control.

**Figure 1.**
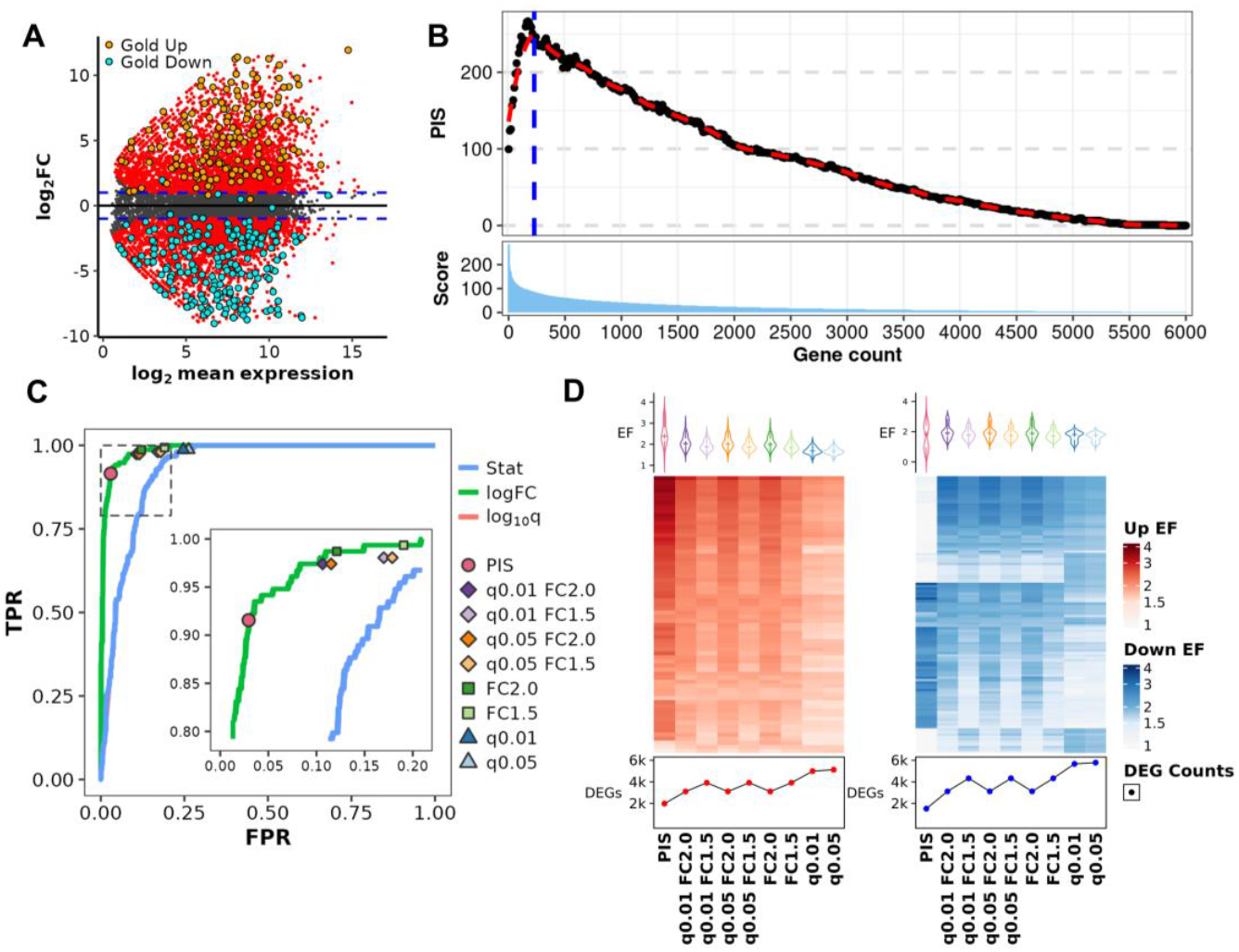
Benchmarking of the Pathway Impact Score (PIS) using the MAQC2 dataset. (A) MA plot illustrating differential gene expression in the MAQC2 dataset. Genes with a fold change (FC) ≥ 2.0 and q-value < 0.01 are highlighted in red. Gold standard DEGs, validated by qRT-PCR, are marked in orange (upregulated) and cyan (downregulated). (B) Demonstration of the PIS peak in the MAQC2 dataset. Genes are ranked based on DESeq2’s stat values and grouped into bins of 10. For each cumulative bin, the sum of pathway enrichment scores is plotted. The red line represents a loess-smoothed curve, with the peak indicating the optimal number of DEGs that maximize pathway-level perturbation signals. This peak pattern is typical for DEG sets ordered by stat values. (C) Comparison of upregulated DEG prediction performance using three gene score metrics: Wald statistics, log2FC, and −log10 q-value. Receiver Operating Characteristic (ROC) curves are presented, with Area Under the Curve (AUC) values indicating predictive accuracy. Sensitivity and specificity for DEGs selected by eight conventional thresholds and PIS are also plotted for comparison. (D) Enrichment Factors (EF) for the top 50 upregulated (left) and downregulated (right) pathways identified by each thresholding method. Higher EF values suggest stronger biological relevance and pathway coherence among the selected DEGs. This figure collectively illustrates that PIS not only aligns well with experimentally validated DEGs but also enhances pathway enrichment, offering a robust and biologically informed approach to DEG selection.

Conventional DEG selection typically relies on predefined combinations of fold change (FC) and false discovery rate (FDR) thresholds. To identify the optimal threshold for PIS, we evaluated pairwise combinations of FC and FDR values. The MAQC2 dataset was chosen due to its controlled design, characterized by minimal inter-replicate variability and a broad distribution of FC values. Thresholds were explored across FC values ranging from 1.2 to 4.0 and FDR values between 0.2 and 0.01, at 0.1 intervals. The DEGs selected under the PIS-optimized thresholds were compared with those obtained using eight commonly applied cut-offs (FDR < 0.01 or 0.05 and FC > 1.5 or 2.0, applied individually or in combination), as well as three gene-ranking metrics from DESeq2: Wald statistic (‘stat’), log2 fold change (‘log2FC’), and signed −log10 FDR. Among these, log2FC yielded the highest performance as a single metric. PIS-selected DEGs closely followed the ROC curve of log2FC but exhibited greater specificity with minimal compromise in sensitivity (Fig. 1C).

The advantage of the PIS approach was further supported by F1 score analyses, which demonstrated superior performance relative to all other threshold combinations (Fig. S1B). Furthermore, pathway enrichment analysis revealed that PIS-optimized DEGs not only recapitulated the major biological processes identified by conventional methods but also achieved higher enrichment scores (Fig. 1D). This is likely attributable of the PIS framework, which intrinsically prioritizes DEG sets that maximize pathway enrichment, thereby enhancing the clarity and strength of pathway-level signals. Consistent results were observed when applying PIS to DEGs identified by edgeR, reaffirming that the method is effective across different DEG analysis tools (Fig. S1C, D).

### Evaluation of Pathway Impact Score method on biological datasets

While the MAQC2 dataset serves as a valuable benchmark, it does not fully capture the complexity and variability inherent in biological datasets, where greater inter-sample variation is common. To assess the applicability of PIS in more realistic biological contexts, we analyzed 14 transcriptome profiles (12 microarrays and 2 RNA-seq) from bleomycin-induced pulmonary fibrosis mouse models, obtained from eight independent studies [30–34]. PIS analysis was performed on each dataset, with DEG thresholds optimized over the same q-value range as used for MAQC2 and a fold change range of 1.2 to 3.0.

When uniform thresholds were applied across all datasets, the number of DEGs varied substantially. In contrast, PIS analysis substantially stabilized DEG counts, highlighting the importance of dataset-specific thresholding (Fig. 2A). Furthermore, pairwise similarity between DEG sets, measured via enrichment factor (EF), was consistently higher when using PIS-selected DEGs compared to those selected by conventional cut-offs (Fig. 2B, Fig. S2).

**Figure 2.**
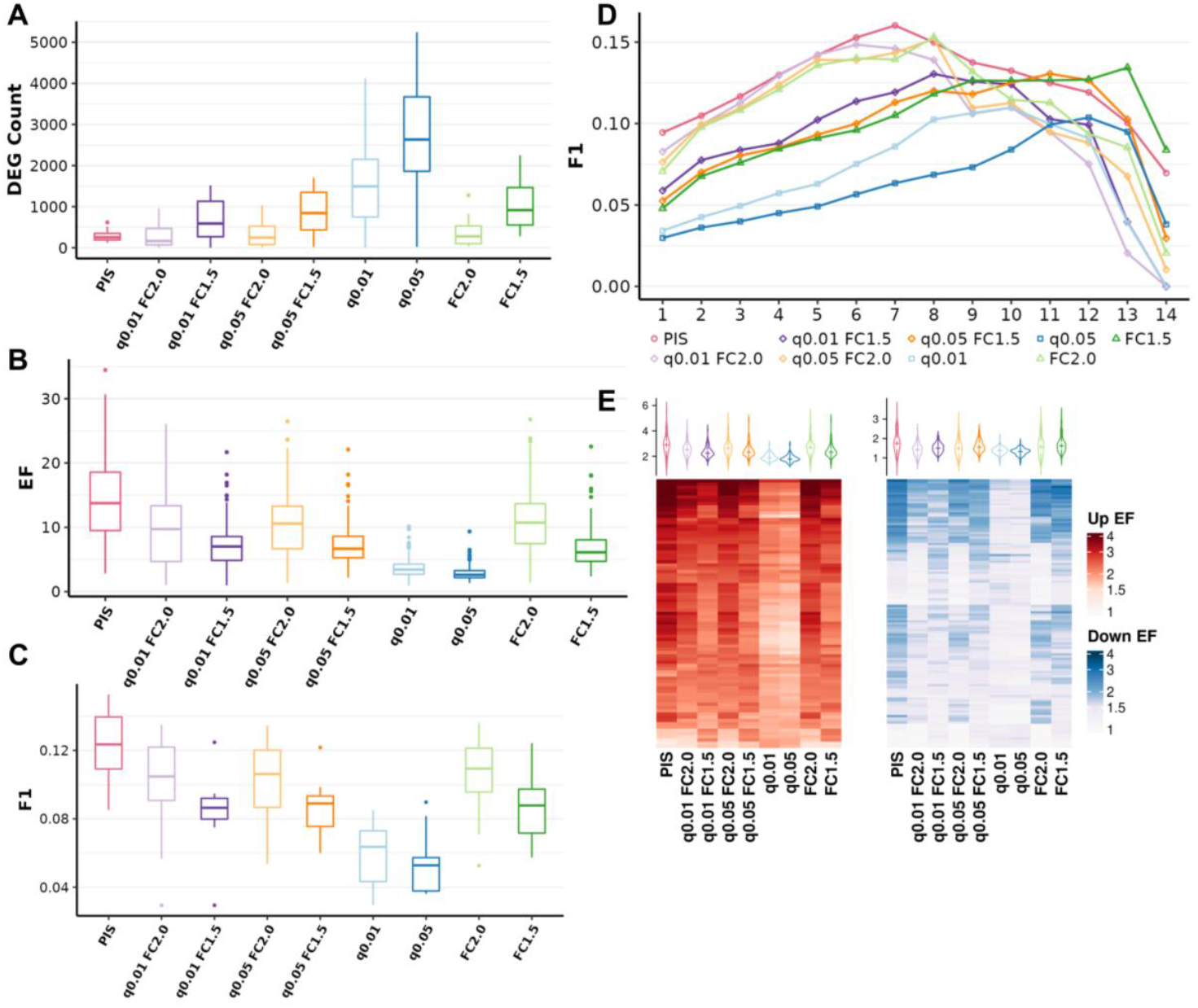
Validation of PIS across pulmonary fibrosis model datasets. PIS consistently enhances DEG reproducibility, biological relevance, and pathway-level interpretability across heterogeneous fibrosis datasets, outperforming conventional thresholds. (A) Total number of DEGs identified by each threshold across 16 public fibrosis transcriptome datasets. PIS yields more conservative but consistent DEG counts compared to conventional cut-offs. (B) Pairwise similarity of DEG sets across datasets, quantified by enrichment factor (EF). PIS shows higher cross-dataset similarity, suggesting improved reproducibility of DEG selection. (C) F1 scores based on DEG set concordance with gold standard genes (identified in at least 8 datasets). PIS achieves the highest overall accuracy in recovering biologically relevant DEGs. (D) F1 scores of consensus genes evaluated under varying consensus thresholds (x-axis: minimum number of datasets a gene must be differentially expressed in). PIS maintains superior performance even at higher stringency, indicating robustness across diverse datasets. (E) Geometric mean of enrichment factors for the top 50 regulated pathways across all datasets. PIS generally shows stronger pathway enrichment, though some downregulated pathways yield lower EF due to its more stringent DEG selection.

To evaluate biological relevance, we compiled a “putative gold standard” list of DEGs associated with pulmonary fibrosis. This reference set was derived from the Human Phenotype Ontology (HPO pulmonary fibrosis) [35], WikiPathways (lung fibrosis gene set) [36], and additional literature reporting genes upregulated in bleomycin-induced fibrosis models [37–39], comprising over 190 genes (Table S3). PIS-selected DEGs demonstrated higher F1 scores relative to conventional threshold methods when benchmarked against this gold standard (Fig. 2C).

Further, we assessed the consistency of DEG selection across multiple datasets by identifying consensus biomarkers—those consistently differentially expressed across varying numbers of datasets. PIS-based thresholds yielded a higher number of consensus genes, particularly as the stringency of the consensus criteria increased (*i*.*e*., requiring presence in more datasets), indicating enhanced reproducibility and robustness in DEG identification (Fig. 2D).

To further assess whether PIS-selected DEGs capture biologically meaningful information more effectively, we compared pathway enrichment scores across different thresholding strategies. PIS-selected DEGs exhibited higher enrichment of activated pathways compared to conventional thresholds (Fig. 2E), despite yielding the lowest number of DEGs. This suggests that PIS identifies a balanced cut-off that preserves strong pathway signals while maintaining a conservative DEG selection. Importantly, pathways implicated in fibrosis pathogenesis— such as inflammation [40,41], extracellular matrix (ECM) organization [42], and fibroblast growth factor signaling [43,44] —were more comprehensively represented in PIS-based analyses (Fig. S3).

The strong agreement of DEGs among the similar datasets, better agreement with known fibrosis genes (‘putative gold standard’), higher number of consensus biomarkers all Collectively suggests PIS’s utility. These findings underscore the importance of accounting for experimental factors, such as replicate number and data quality in DEG selection in the following sections, we further evaluated the robustness of PIS under simulated conditions.

### Effect of Replicate Number on DEG Detection and PIS Robustness

Gene expression datasets are often limited in the number of biological replicates due to practical and cost constraints in experimental design [45]. However, statistical power decreases with smaller sample sizes, reducing the ability to detect true DEGs and increasing the risk of Type II errors [11]. To assess the effect of replicate number, we analyzed two datasets: the MAQC2 dataset and a bleomycin-induced lung fibrosis microarray dataset by Lino Cardenas et al. (GSE34814) [31], consisting of 7 and 5 biological replicates, respectively. Subsampling was performed to generate replicate groups of varying sizes, and differential expression analysis was conducted on each subset.

As expected, applying fixed thresholds led to a marked reduction in the number of DEGs with fewer replicates, with up to 1,000 fewer genes detected between 7 and 2 replicates in the MAQC2 dataset (Fig. 3A). In contrast, DEG counts selected by PIS were less sensitive to replicate number than those identified by conventional methods. Thresholds based solely on fold change (e.g., FC > 2.0, FC > 1.5) exhibited similar stability, as fold change is independent of sample size, and the MAQC2 dataset shows minimal variation by design. Comparable results were obtained using edgeR (Fig. S4) and in the Cardenas dataset, where conventional thresholds frequently failed to detect any DEGs in multiple 2-replicate subsets (Fig. 3B). By contrast, PIS consistently optimized thresholds to yield a reasonable number of DEGs in all but one 2-replicate subset.

**Figure 3.**
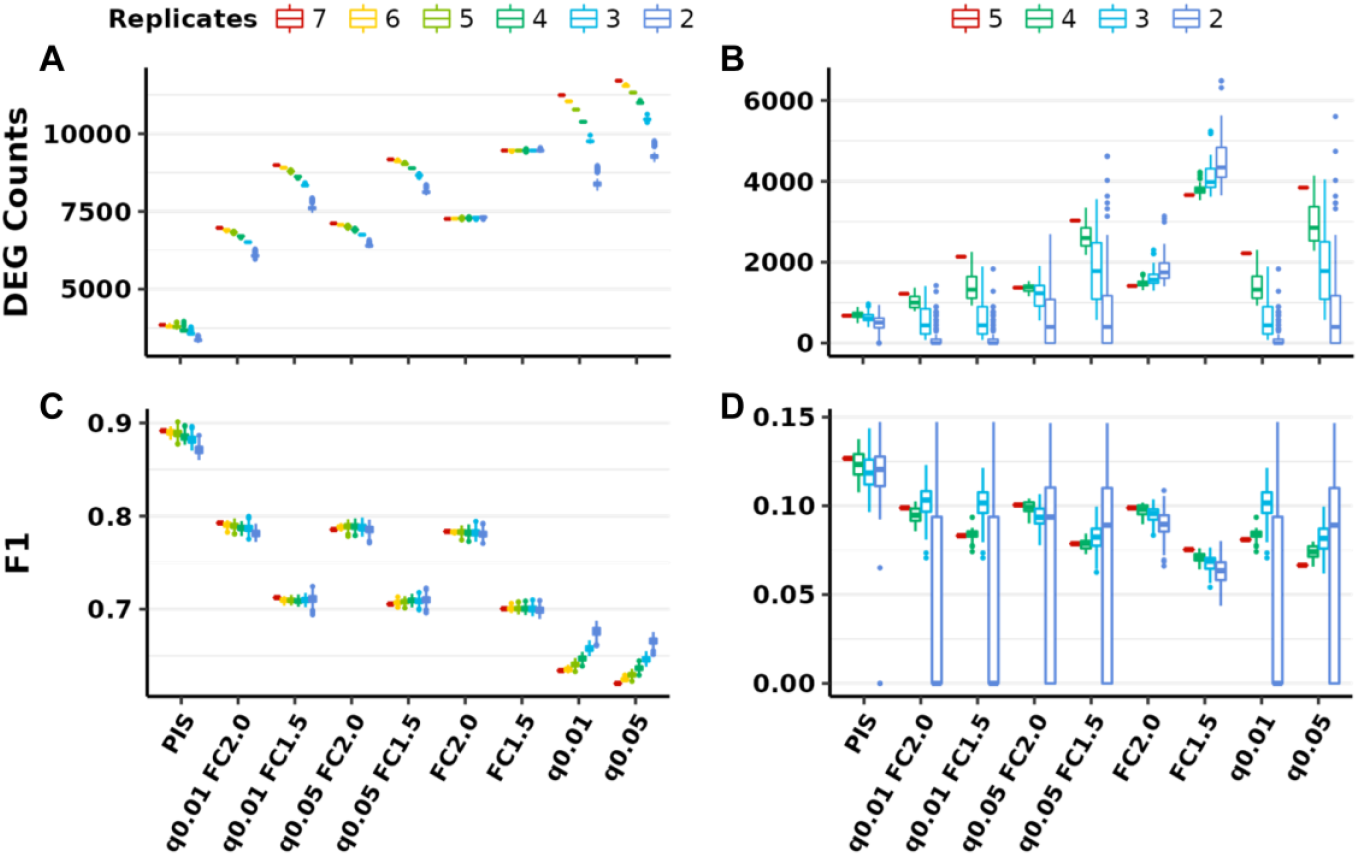
Impact of replicate number on DEG analysis. It tests the robustness of PIS results in comparison with other conventional methods under low-replicate conditions, a common limitation in transcriptome experiments. (A–B) Total number of DEGs identified from replicate subsets of the MAQC2 dataset (A) and the pulmonary fibrosis dataset by Cardenas et al. (B), analyzed using DESeq2. DEG counts generally decrease as the number of replicates decreases. PIS shows more consistent DEG counts across varying replicate numbers compared to conventional thresholds, which show larger fluctuations and often fail to detect DEGs with very small sample sizes. (C–D) Mean F1 scores of DEGs identified from replicate subsets of (C) MAQC2 and (D) Cardenas dataset. PIS maintains higher and more stable F1 scores under decreasing replicate numbers, whereas conventional thresholds show marked performance drops, particularly in the Cardenas dataset with higher biological variability.

We further evaluated performance against gold standard references for each dataset (qRT-PCR validation for MAQC2 and a consensus gene set for Cardenas). In both cases, PIS-selected DEGs achieved higher overall F1 scores compared to those identified by conventional thresholds (Fig. 3C, D; Fig. S4B). Moreover, while the performance of conventional cut-offs varied substantially across replicate subsets, PIS maintained more stable and consistent performance, particularly in the Cardenas dataset, which exhibited greater biological variability.

### Robust DEG Selection Under Simulated Noise

High-throughput sequencing data are inherently prone to noise arising from experimental and technical variability. Such noise, particularly affecting low-expressed genes, can distort statistical measures like fold change, leading to biased DEG identification. To evaluate the robustness of PIS under noisy conditions, we introduced artificial noise by randomly adding or subtracting a fraction of each gene’s average expression from its original read counts (see Methods). Analyses were conducted on both the MAQC2 dataset and a fibrosis mouse model RNA-seq dataset by Liu G. (GSE109913).

In the MAQC2 dataset, DEG counts decreased with increasing noise for all thresholds except those based solely on fold change, which is independent of replicate number and thus less sensitive to sampling variation. Using DESeq2, PIS maintained stable DEG counts across most noise levels, while conventional thresholds showed a sharp decline (Fig. 4A). PIS-selected DEGs more accurately identified true DEGs, as reflected by consistently higher F1 scores (Fig. 4B). Comparable results were obtained with edgeR, confirming the robustness of PIS across different DEG analysis tools (Fig. S5A and S5B).

**Figure 4.**
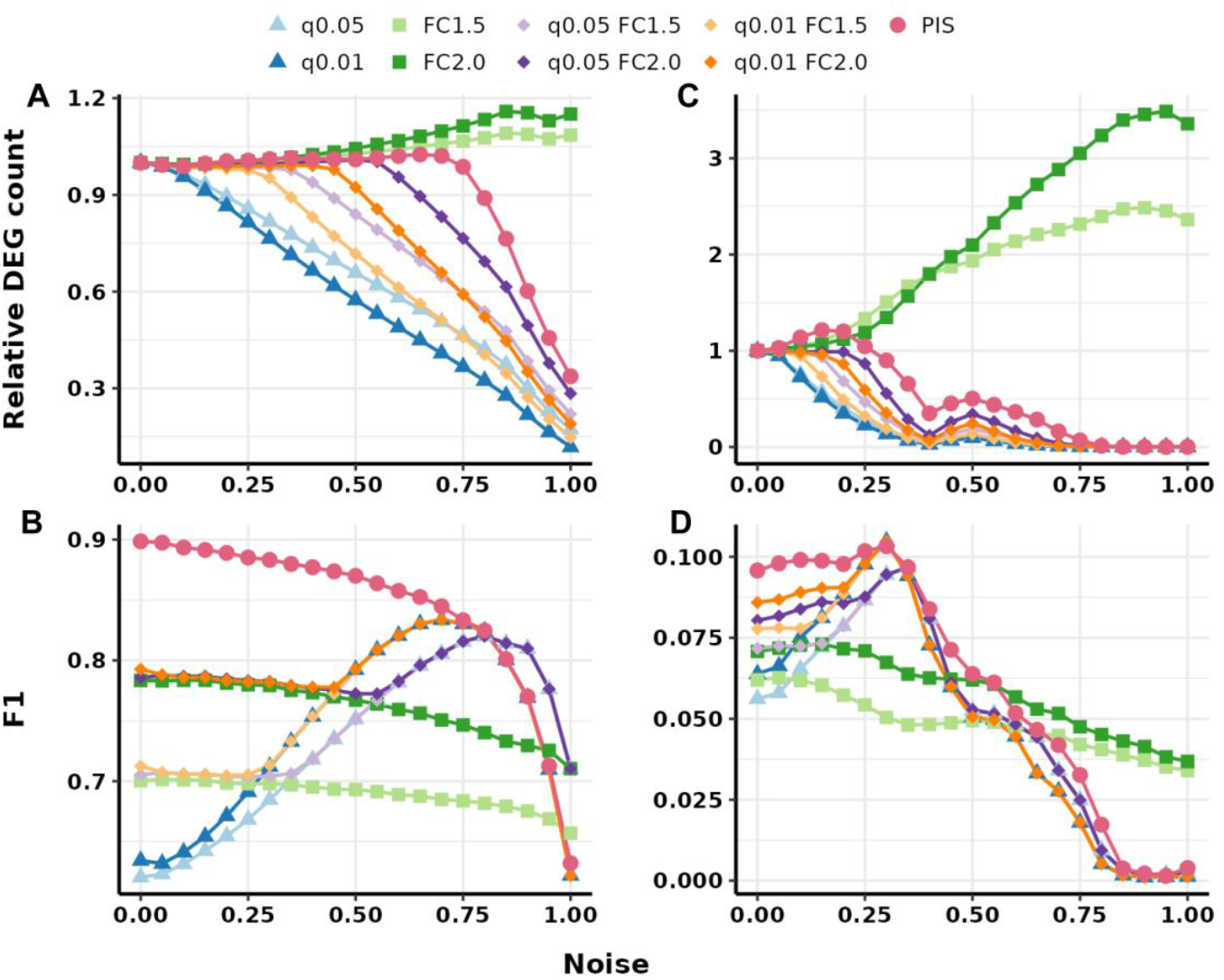
Effect of simulated noise on DEG analysis using DESeq2. It emonstrates the robustness of PIS under low-replicate conditions, a common limitation in transcriptome experiments. (A–B) Total number of DEGs identified from replicate subsets of the MAQC2 dataset (A) and the pulmonary fibrosis dataset by Cardenas et al. (B), analyzed using DESeq2. DEG counts generally decrease as the number of replicates decreases. PIS shows more consistent DEG counts across varying replicate numbers compared to conventional thresholds, which show larger fluctuations and often fail to detect DEGs with very small sample sizes. (C–D) Mean F1 scores of DEGs identified from replicate subsets of MAQC2 (C) and Cardenas (D) datasets. PIS maintains higher and more stable F1 scores under decreasing replicate numbers, whereas conventional thresholds show marked performance drops, particularly in the Cardenas dataset with higher biological variability.

In the Liu G. dataset, which represents a disease model with greater biological variability, DEG counts for all methods declined sharply at 20∼25% noise, and few DEGs were detected beyond this level under conventional thresholds, except for fold change-only criteria. In contrast, PIS thresholds demonstrated higher stability, maintaining reasonable DEG counts even under high noise conditions (Fig. 4C). Moreover, PIS-selected DEGs more effectively captured true fibrosis-related genes, as evidenced by higher F1 scores when benchmarked against the consensus gene set (Fig. 4D). Similar results were observed using edgeR, further supporting the robustness of PIS method (Fig. S5C and S5D).

Taken together, these simulated analyses demonstrate that PIS offers robust and adaptive DEG selection across varying experimental conditions. Whether faced with reduced replicate numbers or elevated noise levels, PIS consistently maintained stable DEG detection and more accurately identified true DEGs compared to conventional thresholds. This consistency was observed across both benchmark (MAQC2) and disease model (Liu G.) datasets, and was independent of the DEG analysis tool employed (DESeq2 or edgeR). By dynamically adjusting threshold criteria in response to dataset-specific variability, PIS preserves biologically meaningful signals, thereby enhancing the reliability of downstream pathway-level analyses across diverse RNA-seq contexts.

### Refining Drug Perturbation Signatures Across Dose Gradients

Pharmacological perturbation often produces dose-dependent transcriptional responses, reflecting differences in target engagement, off-target effects, and cellular adaptation mechanisms. To investigate how PIS accommodates such variability and captures meaningful dose-dependent changes, we profiled transcriptomic responses to four phosphodiesterase (PDE) inhibitors—roflumilast, rolipram, cilostazol, and milrinone—across eleven concentrations spanning nanomolar to micromolar ranges. Notably, these compounds have partially characterized inhibitory potencies (IC50 values) against their primary PDE targets, which allowed us to compare and contextualize dose-dependent transcriptomic responses with known target engagement levels.

As expected, lower doses yielded fewer DEGs due to limited target engagement. PIS, however, adapted thresholds to retain interpretable signatures even at these low doses, unlike conventional fixed thresholds based on fold-change or p-value (e.g., FC > 2.0 or FDR < 0.05), which frequently failed to detect any DEGs, particularly at sub-IC50 concentrations. For example, for cilostazol and milrinone, conventional methods identified almost no DEGs at low to intermediate doses, missing biologically meaningful signals that PIS was able to recover (Supplementary Fig. S6). Importantly, PIS-based DEG counts increased consistently with dose, reflecting stronger perturbation, while conventional methods showed irregular and inconsistent DEG detection across doses, often plateauing or dropping unexpectedly at mid-range concentrations.

We next examined whether PIS could reveal similar transcriptional responses between drugs that target the same enzyme family (Fig. 5). PIS captured clear dose-dependent alignment between PDE4 inhibitors roflumilast and rolipram, with high signature similarity—measured by enrichment factor (EF)—emerging at doses above 125 nM, consistent with their IC50 values. This suggests that transcriptional changes increasingly reflect shared target engagement as the dose increases.

**Figure 5.**
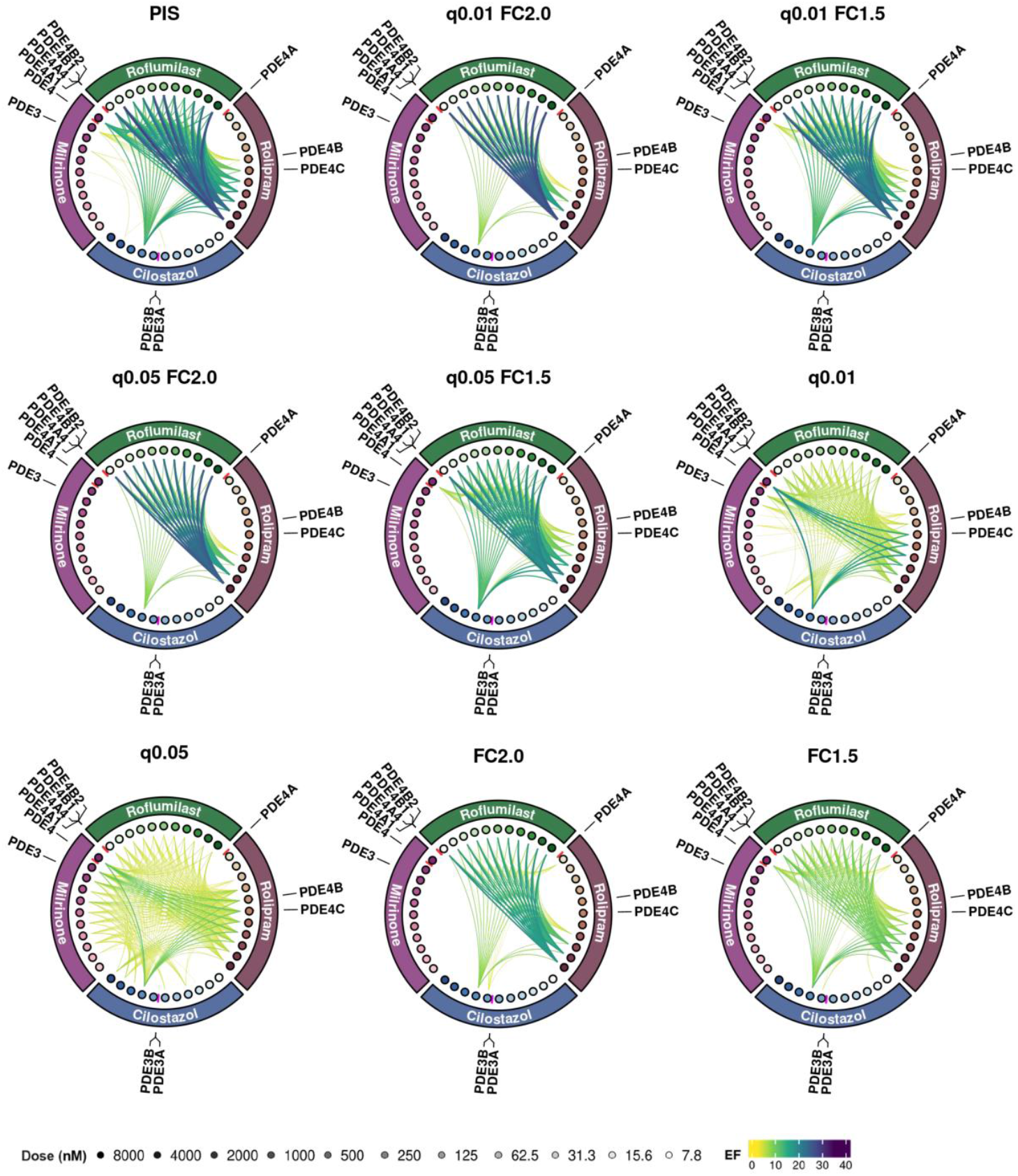
Dose-dependent transcriptomic responses to PDE inhibitors assessed by PIS. Circos plots illustrate the transcriptomic similarity between four phosphodiesterase (PDE) inhibitors— roflumilast, rolipram, cilostazol, and milrinone—across 11 doses (7.8 nM to 8000 nM). Each circle point represents a drug-dose condition, with darker colors indicating higher concentrations. The outer ring text labels correspond to the known PDE isoforms (e.g., PDE3A, PDE4B) targeted by each compound, organized by drug identity and PDE subfamily. Red radial lines adjacent to the target names indicate reported IC_50_ values for each drug–target pair. Curved links represent the top 10% of pairwise DEG signature similarity (EF > 8) between compound-dose pairs, computed by the PIS method. Link color encodes enrichment factor values, ranging from yellow (lower) to deep purple (higher). Greater link density and intensity reflect stronger convergence of transcriptional signatures. PIS-based analysis reveals dose-dependent alignment of transcriptomic profiles, particularly among drugs sharing PDE targets. Notably, higher similarity is observed at pharmacologically relevant concentrations, suggesting that PIS can capture biologically meaningful perturbation patterns aligned with compound potency.

Cilostazol and milrinone, both PDE3 inhibitors, showed fewer DEGs overall, especially for milrinone. However, PIS still detected consistent signature overlap at higher doses near or above their IC50 values. At the highest tested concentration (8000 nM), both compounds exhibited signature overlaps with EF > 8, indicating that even weaker transcriptional responses can converge on meaningful PDE3-related signatures under pharmacologically relevant conditions.

Visual inspection of the Circos plots (Fig. 5) provides a global overview of these relationships. Roflumilast and rolipram displayed dense interconnections at mid-to-high doses, while cilostazol and milrinone showed fewer but distinct links at their highest concentrations. Occasional cross-family connections, such as links between PDE4 and PDE3 inhibitors at higher doses, were also observed, possibly reflecting off-target effects or downstream pathway crosstalk.

Considering the findings from all three validation contexts, the results demonstrate that PIS improves the sensitivity and consistency of DEG detection across dose gradients. It also maintains robustness under artificially introduced noise and enhances the resolution of pharmacological similarity mapping. Notably, similar dose-dependent patterns and compound relationships were observed when applying PIS to DEG results generated using edgeR, supporting the robustness and generalizability of the approach across different DEG analysis methods. By dynamically adapting to perturbation strength and revealing dose-specific and cross-compound transcriptional alignments, PIS is shown to better reflect drug mechanisms of action, optimizing dosing strategies, and supporting comparative pharmacological profiling.

## Discussions

Despite the growing availability of transcriptome datasets, the challenge of defining biologically meaningful and reproducible DEGs remains unresolved. The continued reliance on arbitrary fold change and p-value thresholds contributes to inconsistencies across studies, complicating efforts to compare findings or derive generalizable biological insights. While technical improvements in data harmonization and meta-analysis pipelines have addressed some sources of variability, the scientific arbitrariness of DEG thresholding has received relatively little attention. [46–48].

The Pathway Impact Score (PIS) addresses this limitation by connecting DEG selection to pathway-level biological signals. By identifying thresholds that maximize cumulative pathway enrichment, PIS provides a data-driven alternative to fixed cut-offs. This approach allows DEG selection to be adapted to the specific characteristics of each dataset, improving the consistency of downstream pathway analyses. The results presented in this study demonstrate that PIS can stabilize DEG selection across datasets with varying noise levels and replicate numbers, including both benchmark and biologically complex models.

PIS also offers practical implementation advantages, requiring only standard differential expression results and pathway annotations. Unlike more computationally intensive network-based methods, PIS can be applied with minimal additional modeling or parameter tuning. This simplicity may make it accessible for a range of transcriptome studies, including meta-analyses where dataset-specific variability is a concern. However, it should be noted that PIS relies on the assumption that meaningful biological responses are reflected in coordinated pathway-level changes, which may overlook certain biologically important signals that do not manifest as coordinated pathway-level changes, particularly in poorly annotated systems.

In summary, PIS offers a systematic and biologically informed strategy for deriving data-adaptive DEG thresholds. While its adaptive nature can improve reproducibility and interpretability, it may also introduce bias or alter type I/II error rates, so results should be interpreted with caution. By linking gene selection to pathway-level coherence, PIS can support more consistent transcriptome analyses across diverse experimental designs and biological contexts. Its modest computational demands and reliance on widely available resources make it feasible for both large-scale studies and smaller projects with limited infrastructure. Although its application to single-cell and spatial transcriptomics remains untested, the framework could be adapted to these settings, where technical noise, dropout events, and limited replication pose additional challenges.

## Key Points

1. Pathway Impact Score (PIS) provides a consistent method for selecting optimal DEG thresholds.
2. PIS can maximizes biological interpretation by focusing on pathway perturbations rather than arbitrary cut-offs.
3. This method demonstrated improved accuracy in identifying true DEGs compared to conventional criteria, while being robust to experimental noise and variations in replicate number.
4. PIS can be seamlessly integrated into, or serve as a complementary component of existing transcriptomic analysis pipelines.

## Author Contributions

W.K. designed the study and wrote the manuscript. Y.Y., M.A. and H.L analyzed the data and wrote the manuscript. All authors read and approved the final manuscript.

## Data Availability

The code for PIS analysis is available at https://github.com/littleheroncodes/PIS. The drug-perturbed expression profiles for roflumilast, rolipram, cilostazol, and milrinone are availalbe at GEO (Accession Number : GSE297400)

