## Supplementary Figures and Tables for "Pathway impact analysis (PIS) for robust and comprehensive interpretation of differentially expressed genes (DEGs)"

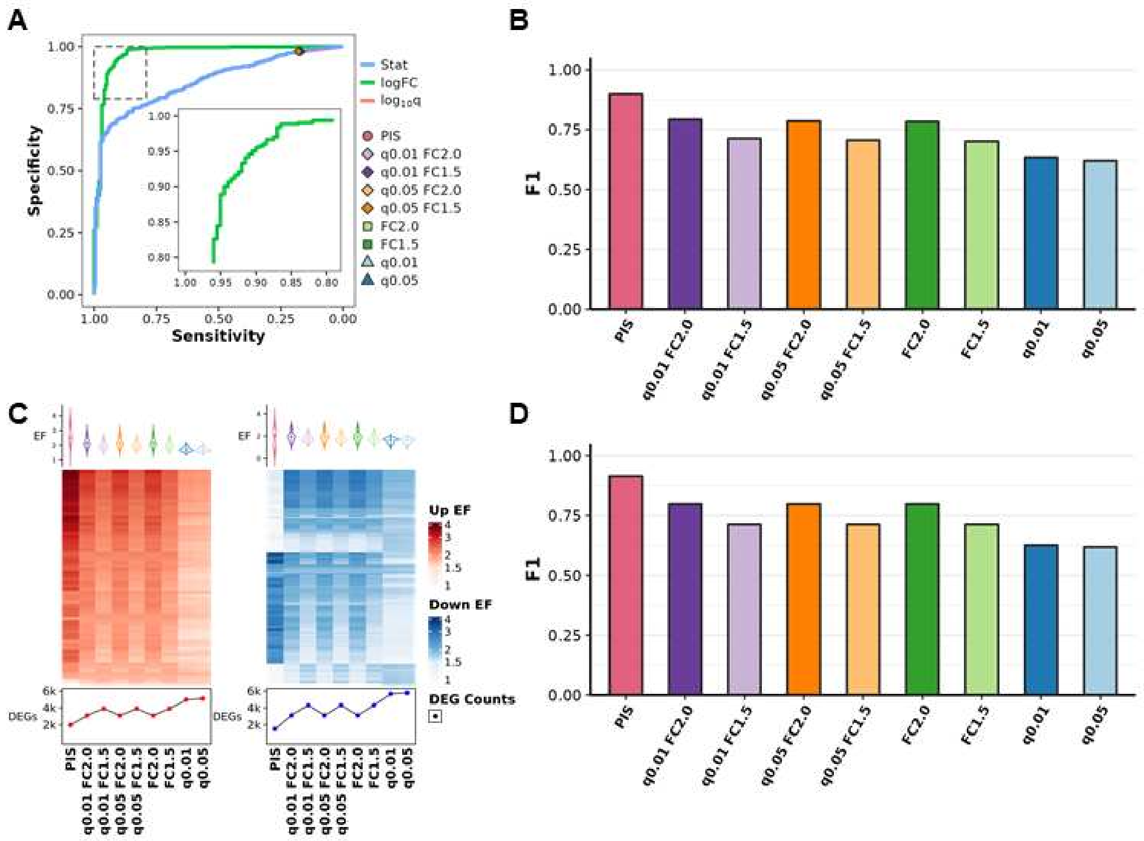


**Supplement Figure S1. Performance evaluation of PIS-selected DEGs. (A)** Down-regulated DEG prediction performance (AUC) of 3 gene score metrics (Wald statistics, log 2FC, log10 q-value). The sensitivity and specificity of DEGs selected by 8 thresholds and PIS are plotted alongside for comparison**. (B)** Median F1 scores for up and down DEGs selected by 8 conventional thresholds using DESeq2 compared to PIS. **(C)** Enrichment Factors of top 50 up (left) or down-regulated (right) pathways selected by each threshold**. (D)** Median F1 scores for up and down DEGs selected by 8 conventional thresholds using edgeR compared to PIS.

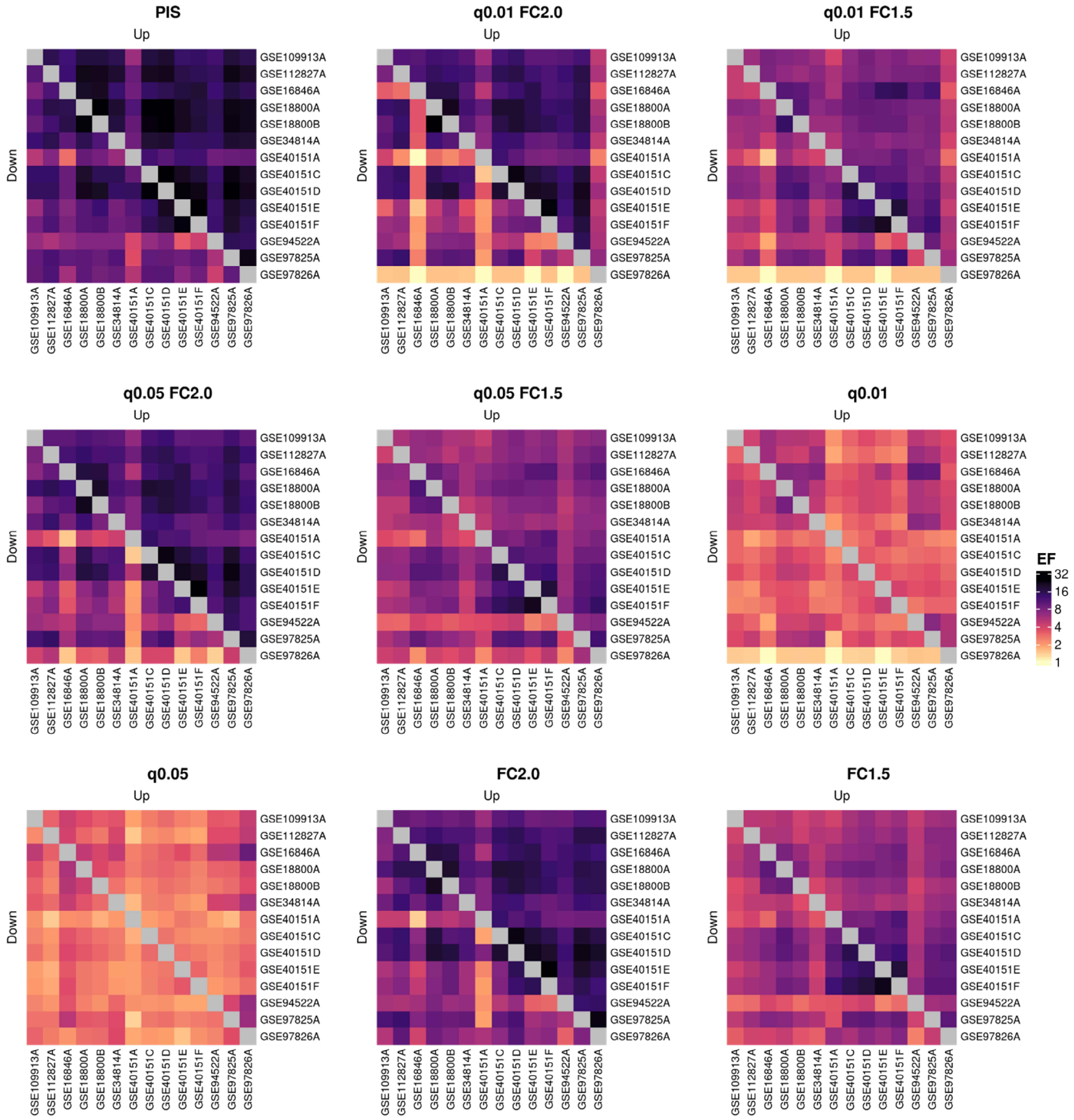


**Supplement Figure S2. DEG signature similarity (EF) between datasets for each threshold.** Enrichment factors between up-regulated DEGs are shown in the upper half, while the lower half of each heatmap shows the EF between down-regulated DEGs.

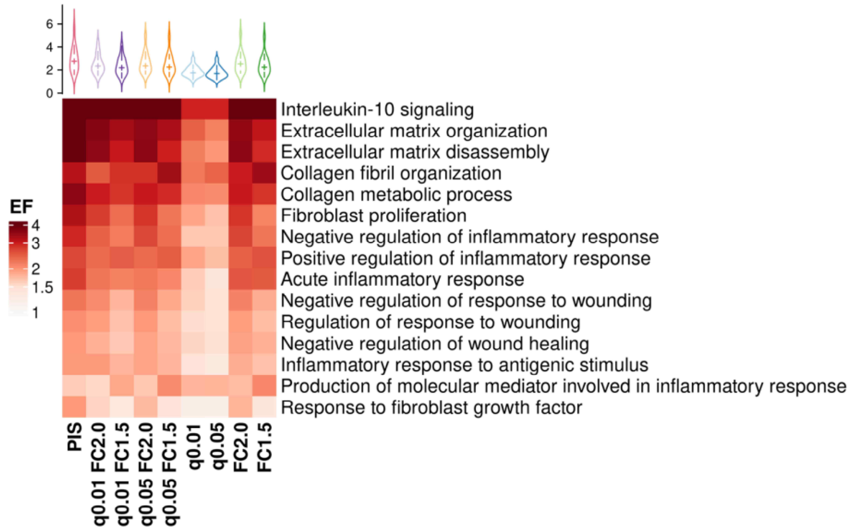


**Supplement Figure S3. Fibrosis-related pathway enrichments scores.** Enrichment of key pathways regulating fibrosis, such as inflammation, extracellular matrix organization. The geometric mean of enrichment factors across 16 profiles was calculated by each threshold for comparison against the PIS method.

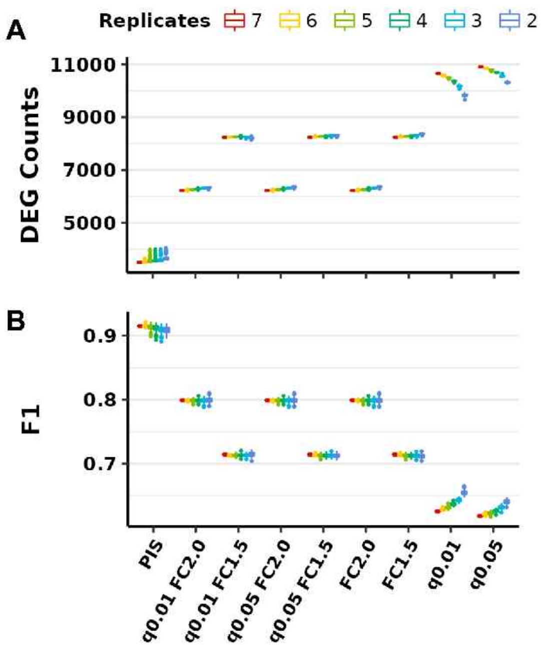


**Supplement Figure S4. Effects of replicate numbers in DEG analysis using edgeR. (A)** The sum of up and down-regulated DEGs selected for replicate subsets in MAQC2 dataset using edgeR. **(B)** Mean F1 scores of up and down DEGs selected for replicate subsets in MAQC2 dataset.

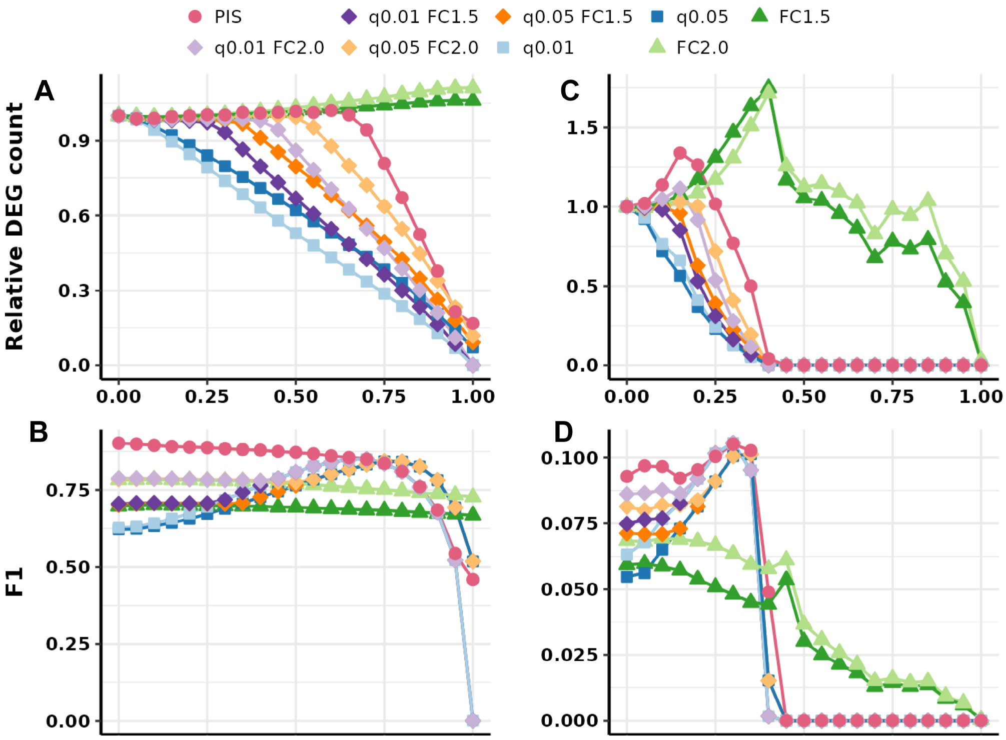


**Supplement Figure S5. Artificially introduced noise effects in DEG analysis using edgeR. (A-B)** Relative mean number of DEGs selected by the level of noise in MAQC2 dataset (A) and lung fibrosis dataset by Liu et al (B). Artificial noise was simulated for 100 iterations and DEG counts were averaged. **(C-D)** DEG performance evaluation by the level of noise in MAQC2 dataset (A) and lung fibrosis dataset by Liu et al (B). F1 scores were calculated for each iteration and averaged.

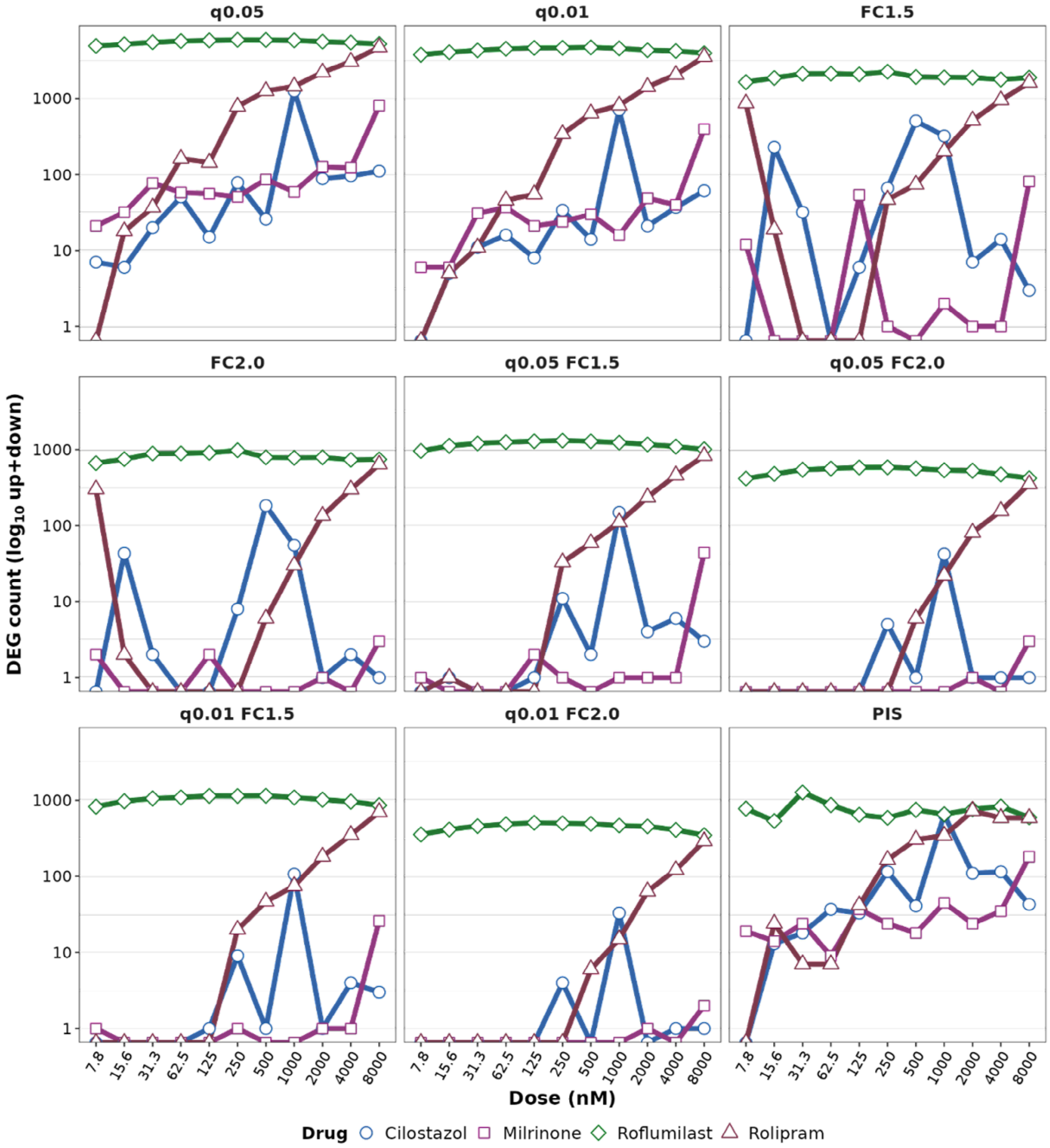


**Supplement Figure S6. Number of DEGs selected for dose-dependent drug treatments.** The sum of up and down DEG counts for 4 PDE inhibitors across 11 doses. The y-axis is in the $\log_{10}$ scale.

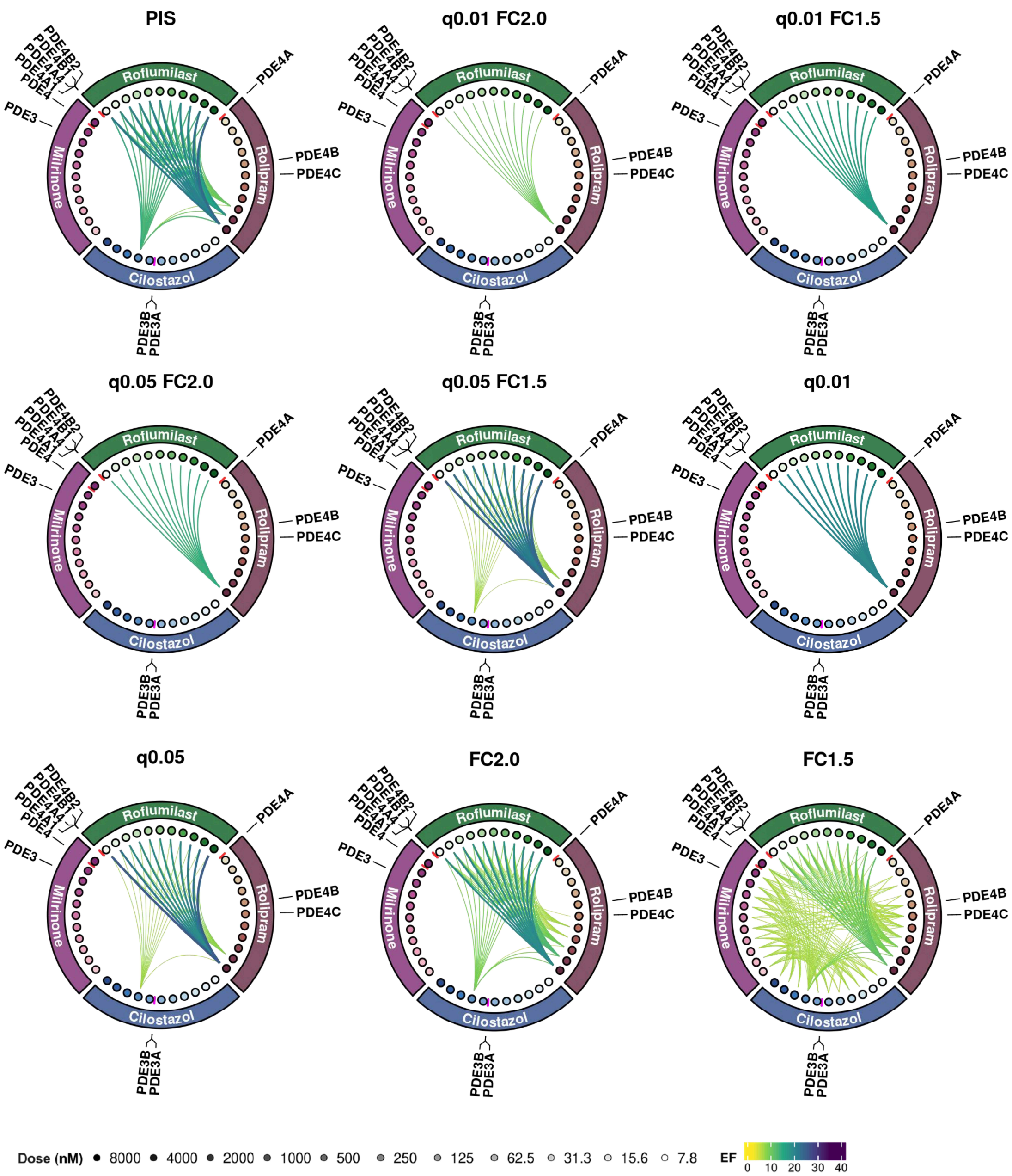


**Supplement Figure S7. Dose-dependent transcriptome analysis of PDE inhibitors through PIS analysis using differential expression results from edgeR.** Circos plot depicting the similarity between transcriptome profiles of 4 PDE inhibitors (Rolipram, Roflumilast, Cilostazol, and Mirlinone) on 11 different doses. Points indicate the dose treatment, with darker colors for higher doses. The IC50 (μM) of each compound against PDE targets are denoted as red lines between the points. DEG similarity (EF) between different compounds where EF > 8 (top 10%) are marked by links.

**Supplementary Table 1 Bleomycin induced pulmonary fibrosis expression data collected from Gene Expression Omnibus (GEO)**

| Analysis ID | GEO Accession | Publication |
|---|---|---|
| GSE109913A | GSE109913 | |
| GSE112827A | GSE112827 | Savary et al. 2019 [26] |
| GSE97826A | GSE97826 | Savary et al. 2019 [26] |
| GSE97825A | GSE97825 | |
| GSE34814A | GSE34814 | Cardenas et al. 2013 [27] |
| GSE94522A | GSE94522 | |
| GSE40151A | GSE40151 | Peng et al. 2013 [28] |
| GSE40151C | | |
| GSE40151D | | |
| GSE40151E | | |
| GSE40151F | | |
| GSE18800A | GSE18800 | Oga et al. 2009 [29] |
| GSE18800B | | |
| GSE16846A | GSE16846 | Scotton et al. 2009 [30] |

**Supplementary Table 2. IC50 of PDE inhibitors**

| Compound | Target | IC50 (nM) | Reference |
|---|---|---|---|
| Roflumilast | PDE4A1 | 0.7 | Hatzelmann et al., 2010 |
| Roflumilast | PDE4A4 | 0.9 | Hatzelmann et al., 2010 |
| Roflumilast | PDE4B1 | 0.7 | Hatzelmann et al., 2010 |
| Roflumilast | PDE4B2 | 0.2 | Hatzelmann et al., 2010 |
| Rolipram | PDE4A | 3 | Mackenzie and Houslay, 2000 |
| Rolipram | PDE4B | 130 | Mackenzie and Houslay, 2000 |
| Rolipram | PDE4C | 240 | Mackenzie and Houslay, 2000 |
| Cilostazol | PDE3A | 375 | Pubchem (mean IC50) |
| Cilostazol | PDE3B | 375 | Pubchem (mean IC50) |
| Milrinone | PDE3 | 2243 | Pubchem (mean IC50) |
| Milrinone | PDE4 | 6000 | Pubchem (mean IC50) |
